# A method of identifying false positives in the strain-specific variant calling of rice

**DOI:** 10.1101/2023.10.02.560417

**Authors:** Sunhee Kim, Sang-Ho Chu, Yong-Jin Park, Chang-Yong Lee

## Abstract

In this study, we investigated the strain-specific effect in genetic variant calling from next-generation sequencing data. For this purpose, we used two major strains of the rice genome, Indica and Japonica, to build different variant calling models that differ in the composition of samples from the two strains. We found that the more the samples differed in their strains from the reference sequence, the more variants were predicted. In particular, the increase in predicted variants was noticeable when the samples that differed in their strains from the reference were included. We used machine learning approaches to understand this finding and compared the performance of different variant calling models using confusion matrices constructed from the predicted variants. We found that a significant proportion of the incrementally predicted variants are potential false positives, which becomes more pronounced the more phylogenetically different accessions from the reference are included in the samples. For the accuracy of the predicted variants, we proposed a method to identify the false positives that can be excluded from the potential false positives if necessary. The proposed method involves calling true variants from the purebred samples. We demonstrated the validity of the proposed method on the different variant calling models and showed a reduction of false positives in the predicted variants. As an example of practical utility, we applied the method to the dbSNP, a database of known variants, and demonstrated a way to identify false positives in the dbSNP. In these respects, this study provides general recommendations for effective practices in strain-specific variant calling in rice.

## Introduction

Advances in next-generation sequencing (NGS) [1] have revolutionized genomic research. It has enabled massively parallel genome sequencing with ultra-high throughput, scalability, and cost-effectiveness. Effective methods, such as those compiled in the Genome Analysis ToolKit (GATK) [2, 3], have contributed to the identification of large numbers of genetic variants, such as single nucleotide polymorphisms (SNPs) and insertion-deletions (indels), across the entire genome within a population. The identified genetic variants are used to test associations with phenotypes of interest, to predict the breeding value of a particular trait or population, and to study genetic diversity within and between populations of species, to name a few [4–6].

Although specific calling pipelines and bioinformatics tools may vary, NGS-based genetic variant calling typically requires a reference sequence, sample sequencing reads in FASTQ format [7], and the dbSNP [8], a database of known variants, for base quality score recalibration. Special attention should be paid to the reference sequence and sample reads if we are interested in predicting genetic variations that are unique to a particular strain (or subspecies) within a population. This is called strain-specific variant calling. A typical example would be variant calling using Indica and Japonica accessions, two major strains of rice (*Oryza sativa*). A large genomic difference between Indica and Japonica [9–11] makes strain-specific variant calling special and important to understand the considerable differentiation between the two strains.

For strain-specific variant calling, it is important that the reference sequence closely matches the samples of a strain or mixture of strains being analyzed. If the reference does not adequately represent the genetic diversity present in the samples, strain-specific genetic variants in the samples may not be detected, resulting in false negatives. If the samples are a mixture of different strains, it may be difficult to detect the presence of strain-specific variants or to distinguish these variants from those shared by different strains. This results in false negatives or false positives, depending on the reference strain. Thus, the presence of multiple strains can increase the complexity of variant calling, potentially leading to increased false positives or false negatives. In short, inadequate sample and reference design for strain-specific variant calling can result in false positives and/or false negatives that affect subsequent analyses. Therefore, it is desirable to systematically investigate the effect of samples from different strains on strain-specific variant calling.

There have been studies of false positives in variant calling from the perspective of bioinformatics and the calling pipeline. Examples include the identification of false positives due to sequencing or alignment errors [12], the use of hard filtering criteria for specific features [13], and the application of different types of bioinformatics tools to variant calling results [14, 15]. These studies investigated the conditions under which variant calling may or may not produce false positive variants, not because of the reference and sample design, but because of the bioinformatics pipeline. When the reference and sample are different strains, the mismatches in the alignment are not only due to genetic variations, but also to the genetic difference in the sequences between them. In addition, since the sample data are usually from a mixture of different strains, it is necessary to understand the extent to which the mixed samples affect false variant calls. In this sense, a quantitative study of strain-specific variant calling can provide a general idea of the sample preparation and handling of the false positive variants.

In this study, we quantitatively investigated the strain-specific effect in variant calling and proposed a method to identify the incorrectly predicted variants. We considered two major rice strains, Indica and Japonica. We found that for a given sample size, more variants are predicted when the samples contain more phylogenetically distinct accessions from the reference strain. Considering that the number of variants called depends on the sample size, this result suggests that sample composition is another factor influencing the variant size. We used machine learning approaches to study the variants by classifying them into two classes: actual (or true) variants and predicted variants. The actual variants were called from purebred samples, while the predicted variants were called from non-purebred samples with different mixtures of the two strains. We built different models, differing only in sample composition, to compare the predicted variants from the models.

The performance of the different models is quantified by the correctness of the predicted variants, using the actual variants as a benchmark. To this end, we introduced the confusion matrix [16] constructed from the actual and predicted variants. We analyzed the confusion matrix using the corresponding metrics in the coding region and in the non-coding region, separately. We showed that the closer the reference and the samples are phylogenetically, the better the performance. More importantly, we found that the more samples containing accessions of the different strains as a reference, the more likely the predicted variants are to be potential false positives.

We proposed a method to identify false positives from the predicted variants and validated the method on the different variant calling models. We found that a significant proportion of the predicted variants were false positives. As a practical example, we applied the proposed method to the dbSNP considering them as predicted variants. We showed that the dbSNP contains non-negligible false positives. This quantitative study of the strain difference between references and samples provides a guide to effective practices for strain-specific variant calling.

## Materials and methods

### Data acquisition and preparation

The data sets used in the study belong to two strains of Indica and Japonica in rice. They are the reference sequences, the dbSNP, and the FASTQ data of purebred and non-purebred samples.

### Reference sequences

The reference sequences used for the variant calling are the ASM465v1 for Indica [17] and the Nipponbare IRGSP-1.0 for Japonica [18]. The ASM465v1 was assembled by the Beijing Genomics Institute, Chinese Academy of Sciences. The IRGSP-1.0, constructed from the genome of a single accession, was assembled from the genome of the Oryza sativa Japonica group from the National Institute of Agrobiological Sciences. The reference sequences of both strains can be downloaded from Ref. [19].

### dbSNP

The dbSNP is a publicly available, intermittently updated repository of known variants, such as SNPs and indels, within and across species. Created from a variety of resources, dbSNP is a collection of VCFs [20], a specified format for storing genetic variants. The Japonica and Indica dbSNPs consist of approximately 25,579,795 and 4,538,989 variants, respectively. The variants are called using ASM465v1 and IRGSP-1.0 for Indica and Japonica dbSNP, respectively. Note that these are the same references we used in this study. These can be downloaded from Ref. [8].

### Sample accessions

We used a core set of 139 cultivated accessions out of 527 accessions from the Korean World Rice Collection (KRICE). KRICE consists of cultivated accessions, collected worldwide by the National Gene Bank of the Rural Development Administration, Republic of Korea. KRICE were classified into three varietal types: landrace, bred, and weedy [21–23]. We determined the ecotypes of the different rice accessions using whole-genome resequencing data [24]. The data were used to construct a circular cladogram tree layout using PHYLIP [25] for the network format and CLC Main Workbench [26] for visualization. The resulting tree was then classified based on the ecotypes of known accessions. The inference of the phylogenetic trees constructed by PHYLIP was used to classify the purebred and the non-purebred accessions, as shown in the Supporting Information S1 Fig.

We used resequencing data generated on an Illumina HiSeq 2500 sequencing system platform. The cultivated KRICE accessions were resequenced, with an average coverage depth of 13.1×. Of 139 cultivated accessions in the core set, we used FASTQ data from

33 purebred Indica accessions and 38 purebred Japonica accessions to call the actual variants. For a fair comparison, we used the same number of purebred accessions for both strains. That is, we randomly selected 33 purebred Japonica accessions out of 38 accessions. We also called predicted variants with different models using different mixtures of Indica and Japonica accessions. We set the number of samples as close to the number of purebred accessions as possible for an unbiased comparison. Thus, we used FATSQ data from 34 non-purebred Indica and 34 non-purebred Japonica accessions to call the predicted variants. In addition, to avoid possible overfitting, we did not use the purebred accessions as members of the samples for the predicted variant calling.

The list of sample accessions for the actual and the predicted variants is provided in the Supporting Information S2 Fig. The FASTQ data are available at https://www.ncbi.nlm.nih.gov/under_BioSample, with the accession number listed in the Supporting Information S2 Fig. The links to all data sets (references, dbSNPs, and samples) are also available at https://github.com/infoLab204/rice_strain.

### Sequence alignment and variant calling

To call genetic variants from NGS data, we used a collection of computational tools available in the Genome Analysis Toolkit (GATK) v4.3.0.0 [27]. A variant calling procedure from the resequencing data using GATK went through a series of processing steps, including data preparation, filtering, mapping, sorting, base quality score recalibration with the dbSNP, and variant calling. The raw data were provided in FASTQ format. BWA-mem2 v2.2.1 [28] and Samtools v1.17 [29] were used to index and align the Japonica reference genome (IRGSP-1.0) or the Indica reference genome (ASM465v1). Duplicate reads aligned at multiple locations were removed and sorted using Samtools v1.17. Next, the raw base quality scores were recalibrated to account for technical errors made by the sequencing machine. Finally, the recalibrated base quality scores were used to detect genetic variants (SNPs and indels). Default settings were used for most of the software and tools used in the analysis. The VCF files containing the genetic variant information were used to construct the confusion matrices and evaluate the performance. These are available at https://github.com/infoLab204/rice_strain.

### Actual and predicted variant callings

Using the GATK pipeline, we called the actual and predicted variants using the strain-specific reference and different sets of samples. The actual variants were called using the purebred samples. For example, the actual variants of Japonica were called using the Japonica reference and 33 purebred Japonica sample accessions. Since both the reference and the purebred accessions are the same strain of Japonica, the variants called should be actual (or true) variants of Japonica. That is, we define the actual variants as the variants called from the reference and purebred accessions of the same strain. The actual variants were classified into coding and non-coding regions obtained from the Japonica and Indica gene information. The same argument applies to calling the actual variants of Indica.

The predicted variants were called using the different calling models, which differ in the preparation of non-purebred samples. We built five different models that have the same sample size but differ in the proportion of samples from the different strains as reference: 0%, 25%, 50%, 75%, and 100%. For example, in the 25% model with the Japonica reference, the samples consist of 25% non-Japonica (i.e., Indica) accessions and 75% Japonica accessions. Note that while the purebred samples were used for the actual variant calling, the non-purebred samples with different proportions of the two strains were used for the predicted variant calling. This allows us to compare the performance of the predicted variants between the different models (or on different sample preparations). Similar to the actual variants, the predicted variants were classified into coding and non-coding regions.

### Confusion matrix and associated metrics

We analyzed the variants called by different models using a confusion matrix, a popular way of analyzing results in machine learning approaches. A confusion matrix is a convenient way to contrast actual and predicted instances, in our case, in a binary 2 *×* 2 matrix. It consists of four different types of outcomes along with the positive (or yes) and negative (or no) classifications. Knowing that the genomic positions of the actual and predicted variants are the locations of the variants in the reference sequence, we classified the predicted variants into four types using the actual variants as a benchmark. If a predicted variant is found at the position (or locus) of an actual variant, it is a true positive (*TP*) at that position in the reference sequence. If not, it is a false positive (*FP*). If a predicted variant is not found at the position of an actual variant, it is a false negative (*FN*). If neither a predicted nor an actual variant is found at a position, it is a true negative (*TN*).

Note that the set of actual variants called from purebred samples is not complete in the sense that the purebred samples used are not large enough to call a sufficient number of actual variants. Nevertheless, we can use the set of actual variants as a benchmark for the accuracy of the predicted variants for the following reasons. The confusion matrix is used to compare the results of variant calls from different prediction models. A set of predicted variants from each model is examined against the set of the same actual variants. That is, the results of different models are compared against each other while the set of actual variants is fixed. This allows us to impartially compare the performance of different models as long as we are using the same actual variants.

With the confusion matrices, we can compare the performance of different models using appropriate metrics. One of the simplest metrics is accuracy, which is the ratio of correctly predicted instances to total instances. Although accuracy is a commonly used metric, it can be misleading when used with unbalanced data, where instances of different classes are not balanced. This is because the accuracy does not distinguish between correctly classified instances of different classes. It is known that genetic variants occur approximately every 1,000 nucleotides or about 0.1% of the total genome [30]. Thus, the instances of variant and non-variant are highly unbalanced. In addition, we are interested in instances that belong to the positive class for either actual or predicted variants.

To better measure the performance of unbalanced data, we used three metrics: precision, recall, and F1 score. Both precision and recall are critical when the positive class (i.e., variant) is more important relative to the negative class (i.e., non-variant). Precision and recall are defined as

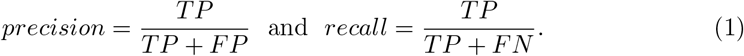

Precision quantifies the ratio of true positives to all positive predictions, or how often a prediction is correct when it predicts a positive. The higher the precision, the lower the false positives. Recall quantifies the ratio of predicted positives to total positives, or how often a prediction is correct when it is actually positive. The higher the recall, the fewer the false negatives. The F1 score is the harmonic mean of precision and recall. It takes into account both false positives and false negatives. Therefore, it performs well on an unbalanced data set and is useful when we need to consider both precision and recall at the same time. It is defined as

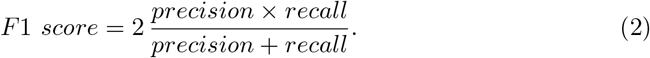

## Results and Discussion

### Model performance comparison

We discussed the results of variant calling with the Japonica reference in detail. We obtained similar results with the Indica reference, which will be briefly discussed later. We called variants using different models (i.e., different sample types) of the same sample size and analyzed variants in coding and non-coding regions separately.

To begin with, we plotted the number of variants called by the different models using the Japonica reference, as shown in Fig. 1. Of the total variants, about 5% variants are found in the coding region and the rest are found in the non-coding regions, regardless of the variant calling models. From Fig. 1, we find that the non-purebred samples predicted more variants than the purebred samples, both in the coding region and overall. Moreover, the higher the proportion of non-Japonica (i.e., Indica) accessions in the samples, the more variants were called. In particular, the incremental number of variants is pronounced when non-Japonica accessions are included in the samples (e.g., samples with 25% non-Japonica accessions). Considering that the different models have the same sample size, these results are inconsistent with the general expectation that the number of variants called is closely related to the sample size. Similar characteristics were found in the non-coding region, as shown in the Supporting Information S3 Fig.

**Fig 1.**
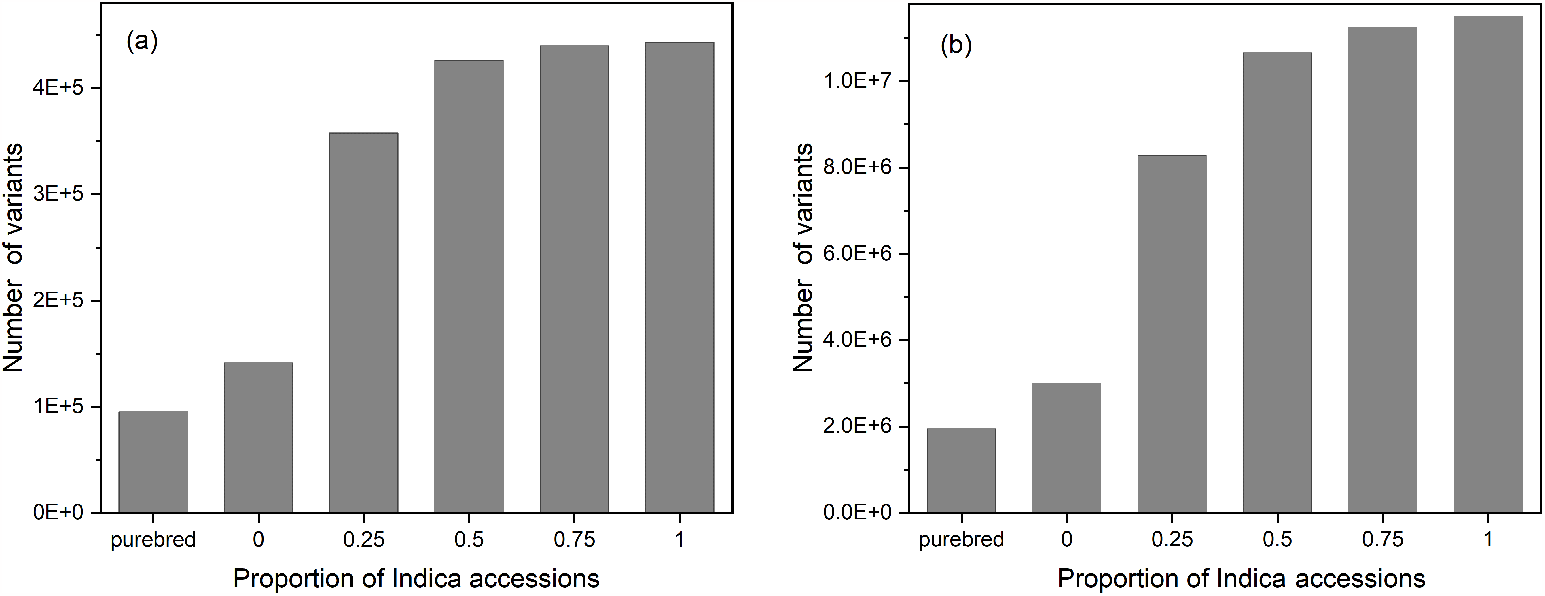
The number of variants called from different models using the Japonica reference (a) in the coding region and (b) overall. Note that the horizontal axis represents the proportion of non-Japonica (i.e., Indica) accessions in the samples, except purebred Japonica.

To understand the unexpected result, we constructed confusion matrices to evaluate the performance of all five models using metrics such as precision, recall, and F1 score. As Fig. 2 shows, while recall is about the same for all models, the precision decreases significantly once non-Japonica accessions are included in the samples. This significant decrease in precision is closely related to the sudden increase in the number of variants shown in Fig. 1. Because of the decrease in precision, the F1 score also decreases, making the overall accuracy worse than the Japonica-only model (i.e., the model with zero proportion of Indica accessions). As shown in Fig. 2, the F1 score of the other models is only half that of the Japonica-only model. Thus, the Japonica-only model outperforms the others, implying that samples should be the same strain as the reference as much as possible for better performance. Similar results were found in the non-coding region, as shown in the Supporting Information S4 Fig.

**Fig 2.**
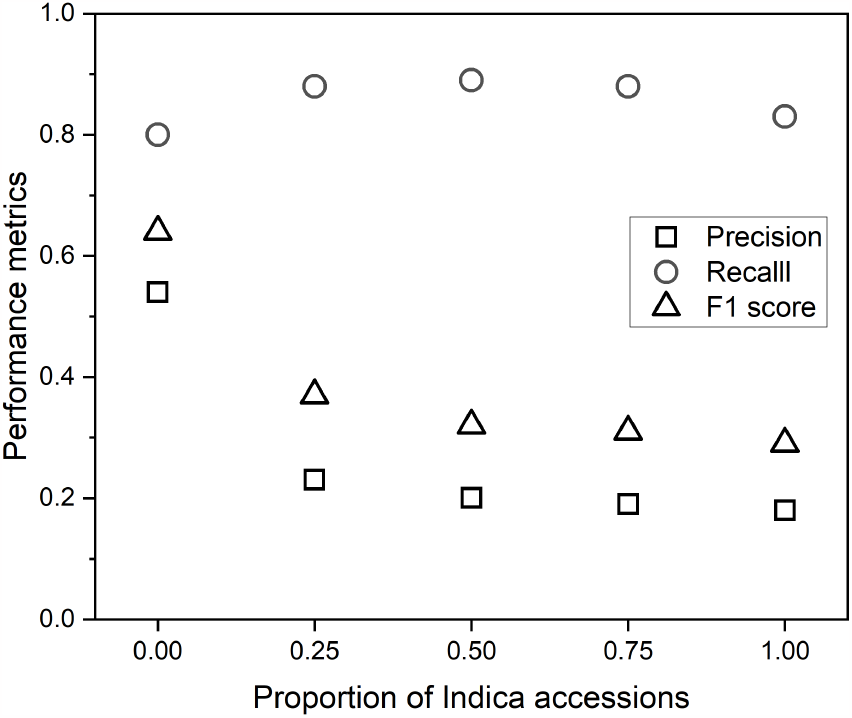
A plot of the performance metrics of precision, recall, and F1 score for different models in the coding region using the Japonica reference. Note that the horizontal axis represents the proportion of non-Japonica (i.e., Indica) accessions in the samples.

According to Eq. (1), a decrease in precision means an increase in the number of false positive variants. To investigate an increase in false positives more closely, we

compare the confusion matrices of the two models: the Japonica-only model and the 25% Indica-mixed model. As shown in Fig. 3, the Japonica-only model had about 10% lower recall (R=0.80) than the other model (R=0.88), which meant that the

**Fig 3.**
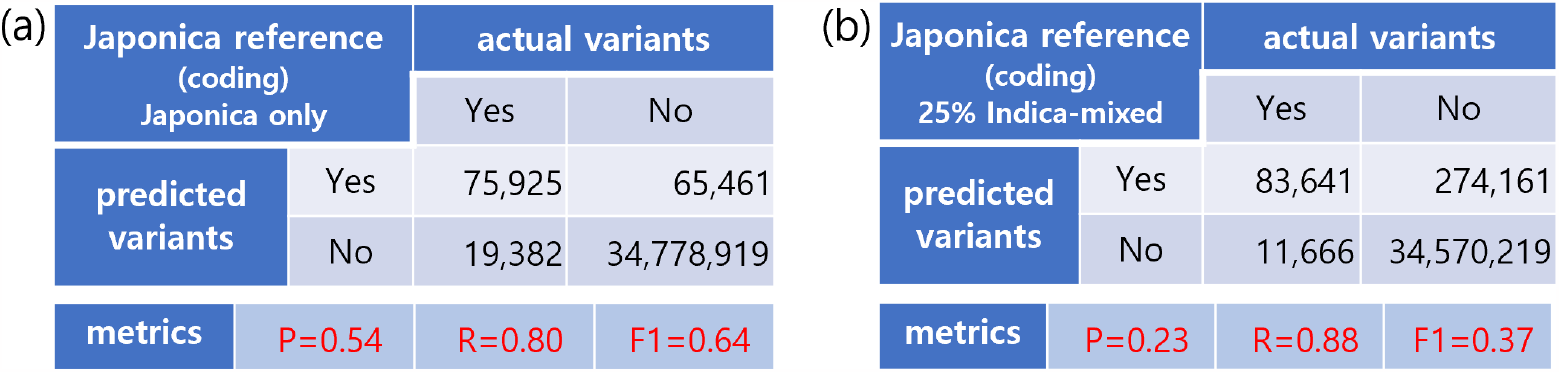
Results of the confusion matrices and their metrics constructed from variant calling in the coding region using the Japonica reference from (a) the Japonica-only model and (b) the 25% Indica-mixed model.

Japonica-only model predicted about 10% fewer actual variants. In contrast, the 25% Indica-mixed model had an estimated precision of P=0.23 as shown in Fig. 3(b), which is much lower than P=0.54 of the Japonica-only model. This indicates that about 77% of the predicted variants from the 25% Indica-mixed model were potential false positives, which is more than four times as many as from the Japonica-only model (274,161 vs 65,461). We used the term “potential” for false positives because not only the false positives, but also new and/or known Japonica variants may be included. In addition, the 25% Indica-mixed model had the highest precision among all Indica-mixed models. This means that other models produce more potential false positives than the 25% Indica-mixed model, as shown in Table 1. Note that the variants identified from the Japonica-only model with the Japonica reference are not necessarily Japonica variants as precision (P=0.54) shows in Fig. 3(a). This is because the samples in the Japonica-only model are not purebred. That is, the Japonica-only model also produces potential false positives, but much fewer in number than the Indica-mixed models. We performed the same type of analyses for the non-coding region and found similar characteristics as shown in the Supporting Information S5 Fig and S1 Table.

**Table 1.** The number of potential false positives and precision from the models of different Indica-mixed proportions.

| Model | 0% | 25% | 50% | 75% | 100% |
| --- | --- | --- | --- | --- | --- |
| Number of potential false positives | 65,461 | 274,161 | 341,829 | 356,560 | 364,265 |
| Precision | 0.54 | 0.23 | 0.20 | 0.19 | 0.18 |

We suggest that the unexpected potential false positives are likely due to the large differences in genome structure and gene content between the two strains. Sample sequences from a strain that differs from the reference sequence are more likely to mismatch with the reference than those from the same strain. This means that the more the samples differ in their strains from the reference, the more false positive variants will be called.

### Identification of false positives

The potential false positives using the Japonica reference consist of the Japonica variants and non-Japonica variants (i.e., the false positives). The Japonica variants, in turn, consist of new Japonica variants and known Japonica variants. The “known” here means variants that have been identified but unrecognized, for example, due to an incompleteness of the actual variants.

To decide whether a potential false positive was either a false positive or a variant, we used information about the strain of the accessions from which a variant was called. The VCF file contains such information. The VCF is the variant call format for storing sequence variations. Each row of VCF represents a variant, and the columns contain attributes about the variants. Starting with the 10th column, additional information is given about the accessions in the samples. The additional information includes the genotype, which indicates the zygosity at the location of the variant. Zygosity allows us to identify the accessions from which a variant was called. Since the strain information of an accession is known, we can identify the accessions and their strains from which the variant was called.

Using the accessions and their strains of variants, we proposed a method to construct a set of false positives, which we called dbFP, meaning the database of false positives.

We explain how to construct the Japonica dbFP, which is a set of false positives with the Japonica reference. We perform variant calling from the samples of both purebred Japonica and purebred Indica accessions with the Japonica reference sequence. The resulting variants can be categorized into three types: variants called from purebred Japonica accessions only, shared variants called from both purebred Japonica and purebred Indica accessions, and variants called from purebred Indica accessions only. The results of categorized variants with the Japonica reference are shown in Fig. 4.

**Fig 4.**
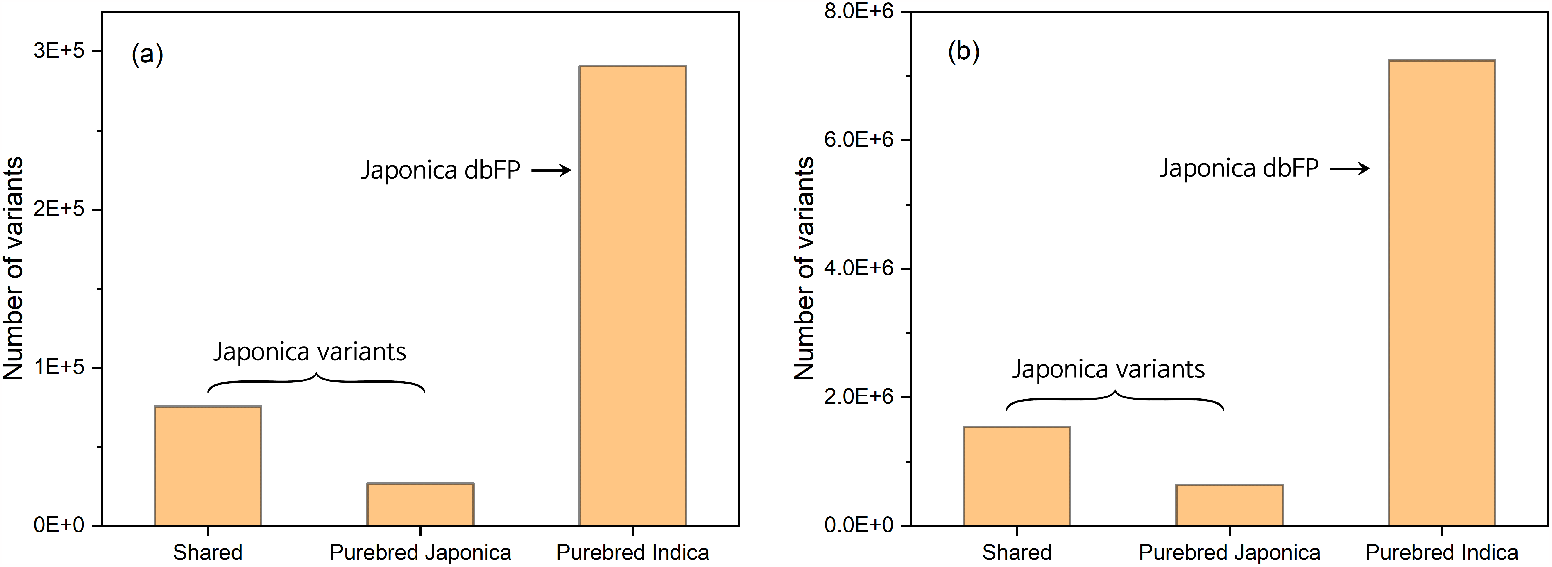
The number of variants called from three different types of samples (a) in the coding region and (b) overall. The “shared” means variants called from both purebred Japonica and purebred Indica accessions.

Since the samples consist only of purebred accessions, the first type should be the Japonica variants. This follows from the definition of the Japonica variant: the variants from the Japonica reference and the purebred Japonica samples are Japonica variants. The variants of the second type are also Japonica variants because the samples contain purebred Japonica accessions. Even if the purebred Indica accessions are removed from the VCF file, the mismatches with the reference are variants because of the remaining purebred Japonica accessions. Therefore, they are also Japonica variants. Variants of the third type are only called from purebred Indica samples. It is not sure if they are Indica variants because we use the Japonica reference, not the Indica reference.

However, they are certainly not called from purebred Japonica samples. So they are not the Japonica variants and are false positives from a Japonica strain perspective. Thus, the variants called from only purebred Indica accessions belong to the Japonica dbFP, as indicated by an arrow in Fig. 4. The Indica dbFP can be constructed in a similar way. Note that the above argument is at least theoretically valid, if not practically so, in the sense that such a constructed Japonica dbFP does not contain all possible Japonica variants. The more purebred samples are used in variant calling, the more likely the argument is to be practically valid. Similar to the set of actual variants, the set of false positives becomes more complete as more purebred samples are used in variant calling.

To demonstrate the usefulness of the dbFP, we considered two confusion matrices constructed in the coding region with the Japonica reference. Figure 5(a) shows the confusion matrix with the 50% Indica-mixed samples from the two strains, and Fig. 5(b) shows the confusion matrix with the same samples as in Fig. 5(a), except that the potential false positives listed in the Japonica dbFP are excluded. Compared to

**Fig 5.**
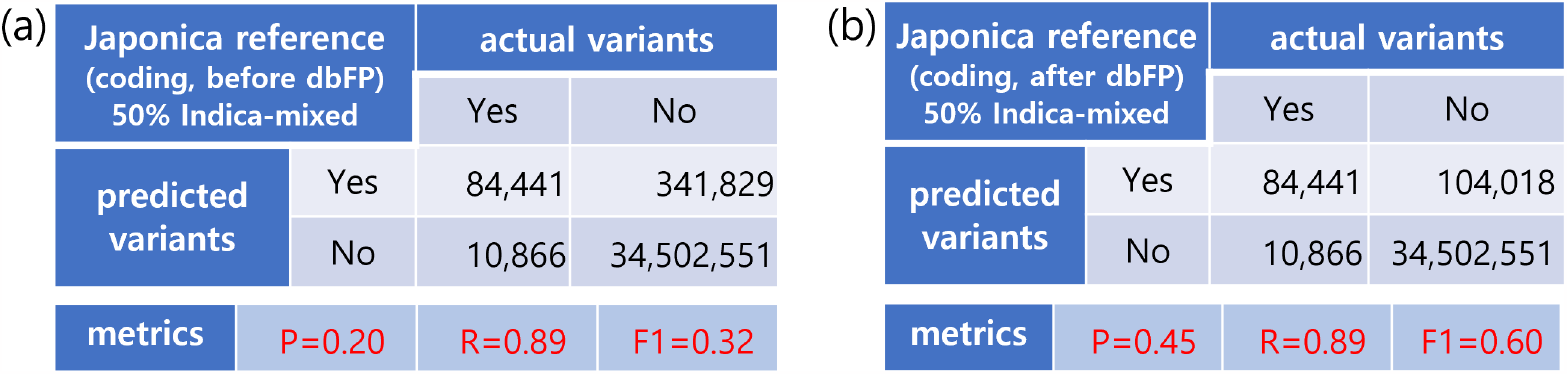
Results of the confusion matrices and their metrics constructed from variant calling in the coding region with the Japonica reference using (a) the half-mixed samples from both strains and (b) the same as (a) except that the variants in the Japonica dbFP are excluded.

Fig. 5(a), we can see from Fig. 5(b) that the precision increases more than double (from P=0.20 to P=0.45), implying a significant reduction in potential false positives (from 341,829 to 104,018). Numerically, the 341,829 potential false positives can be divided into 237,811 false positives identified by the dbFP, and the remaining 104,018 variants including newly found Japonica variants and/or known Japonica variants. We also found similar characteristics in the non-coding region, as provided in the Supporting Information S6 Fig.

The proposed method identifies as many false positives as possible while leaving the true variants intact. The proposed method may not identify all false positives due to the incompleteness of the purebred samples. However, it does not misidentify the true variants as false positives. We expect the proposed method to become more effective and accurate as more purebred samples are used in variant calling. In this respect, dbFP can play a role in the database of false positive variants as dbSNP does in the database of true variants. The dbFP has two columns (chromosome number and the reference position) and can be downloaded from

https://github.com/infoLab204/rice_strain/blob/main/results/dbfp_japonica.gz.

We also performed variant calling with the Indica reference using the same models as with the Japonica reference, except for the exchange of strains in the samples. For the variants called by different models with the Indica reference, we performed the same analyses as with the Japonica reference. Although the numerical details of the analysis results with the Indica reference were different from those with the Japonica reference, the features and trends of the analysis results showed the same characteristics as in the case of the Japonica reference. The analysis results with the Indica reference are given in the Supporting Information S1 File.

### Applying the method to dbSNP

As a practical application, we applied the proposed method to the dbSNP of Japonica and identified the false positives in the Japonica dbSNP. The dbSNP is a publicly available collection of known variants in the form of VCF, a specified format for storing genetic variants. Studies on the quality of the dbSNP revealed high false positive rates due to errors in base calls and bioinformatic tools [31, 32]. We approached dbSNP quality from a strain perspective and investigated whether the Japonica dbSNP contains non-Japonica variants that are false positives. For this purpose, we have introduced the error rate of a sample. It is defined as the ratio of the number of mismatched bases with references not listed in the dbSNP to the total number of mismatched bases in a sample. It is also indirectly related to the recalibration of the base quality score. Note that mismatched bases were counted based on reads, not samples. Therefore, there may be multiple mismatches at the same locus. The more mismatched bases are listed in the dbSNP, the lower the error rate. When using the Japonica dbSNP, the error rate of Japonica samples is expected to be lower than that of non-Japonica (or Indica) samples, because the Japonica dbSNP should by definition contain Japonica variants.

We evaluated the error rates of four different types of same-size samples: purebred Japonica, non-purebred Japonica, purebred Indica, and non-purebred Indica. As shown in Fig. 6(a), we found that the error rate of Indica samples, whether purebred or not, is lower than that of Japonica samples. This means that the mismatched bases of Indica samples are more likely to be listed in the Japonica dbSNP than those of Japonica samples. This result is contrary to expectations. A plausible explanation for this unexpected result is that the Japonica dbSNP includes not only the Japonica variants but also the non-Japonica variants, whose frequency is non-negligible.

**Fig 6.**
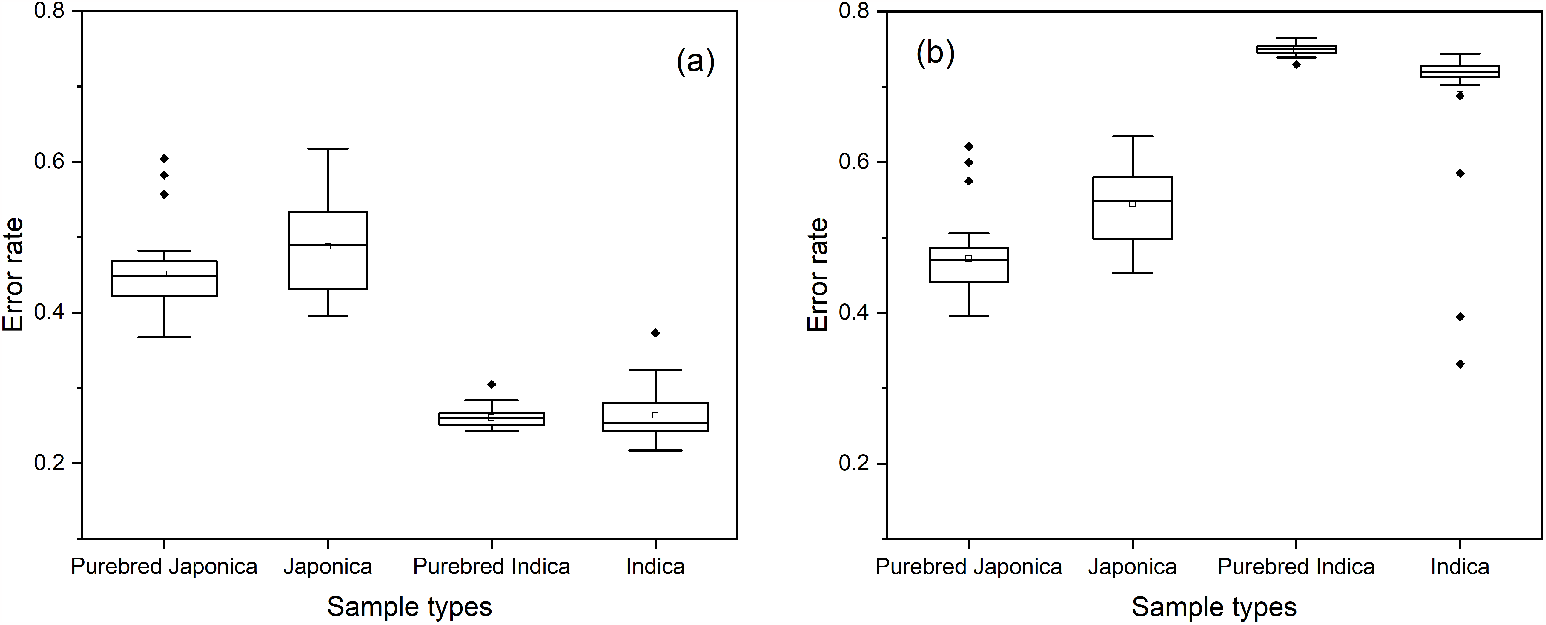
Box plots of the estimated error rates of the purebred and non-purebred Japonica samples and those of the Indica samples (a) before and (b) after exclusion of the false positives from the Japonica dbSNP.

To validate the above claim, we counted the variants shared between Japonica dbSNP and Japonica dbFP, and represented the logical relationships between the two sets of variants using the Venn diagram. Fig. 7(a) shows the result, from which we find that about 20% of the variants in the Japonica dbSNP overlap with the Japonica dbFP. We call this overlap the shared set of variants. This result means that about 20% of variants in the Japonica dbSNP are not Japonica variants, but false positives. The Japonica dbSNP contains non-negligible non-Japonica variants, and the Japonica dbFP plays a role in identifying the false positives (i.e., non-Japonica variants) in the Japonica dbSNP. Once we use the Japonica dbFP to exclude the false positives from the Japonica dbSNP, we have the corrected Japonica dbSNP, although not completely due to the incompleteness of the Japonica dbFP.

**Fig 7.**
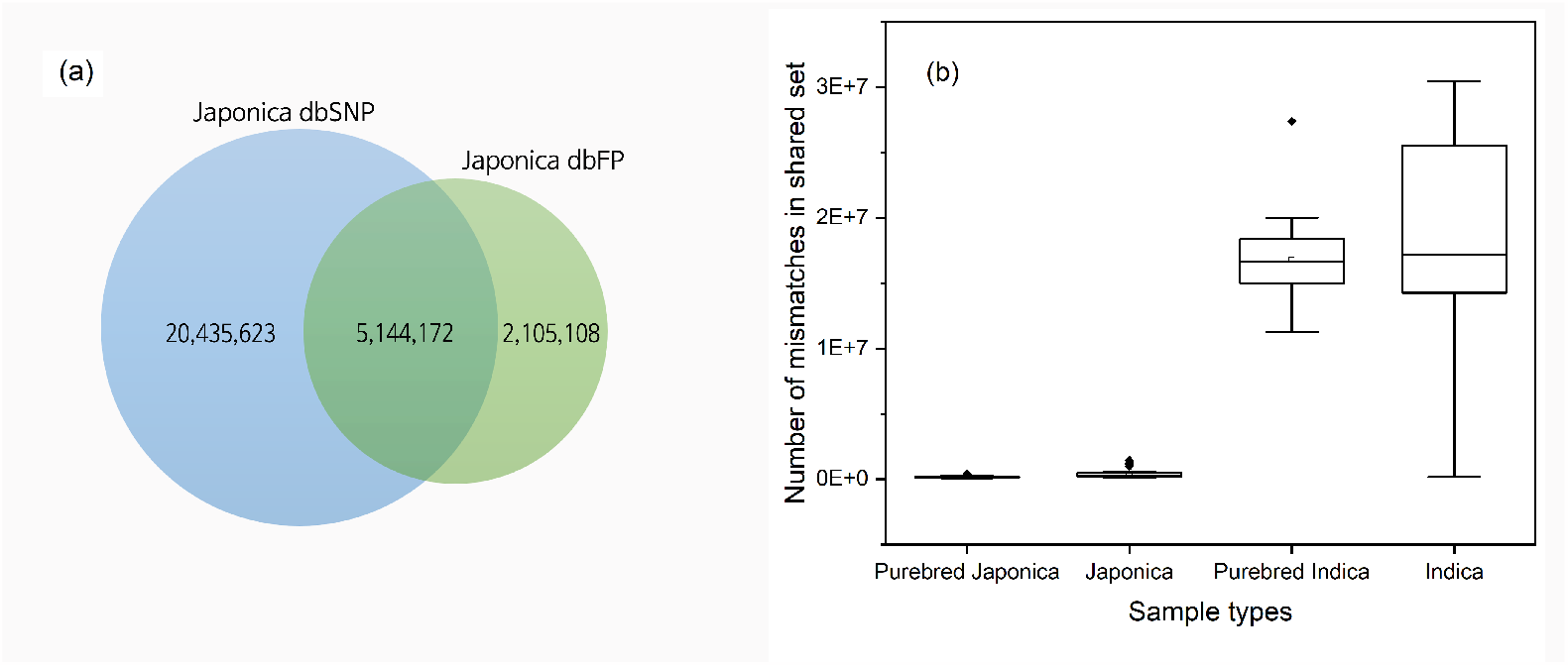
(a) A Venn diagram drawn from the Japonica dbSNP and the Japonica dbFP. The numbers in the diagram are the number of variants in each area. (b) Box plots of the number of mismatched bases in a sample listed in the shared set of variants for each sample type.

It can be assumed that the lower error rate of the Indica samples shown in Fig. 6(a) is due to non-negligible amounts of non-Japonica variants in the shared set. To verify this assumption, we counted the number of mismatched bases listed in the shared set of variants. As shown in Fig. 7(b), Indica samples have far more mismatched bases listed in the shared set than Japonica samples. This implies that the mismatched bases of Indica samples are less likely to be considered errors than those of Japonica samples according to the Japonica dbSNP. This explains the lower error rate of Indica samples than Japonica samples shown in Fig. 6(a).

We re-estimated the error rate using the corrected Japonica dbSNP, and the result is shown in Fig. 6(b), from which we can see that the error rate of the Japonica samples is lower than that of the Indica samples, as expected. The error rate of the Japonica samples has hardly changed, while the error rate of the Indica samples has become much higher than before. This indicates that non-Japonica variants are mostly excluded in the Japonica dbSNP. Thus, the corrected dbSNP contains less number of non-Japonica variants than the original dbSNP, as desired. We obtained a similar result with the dbSNP of Indica as provided in the Supporting Information S1 File.

## Conclusion

In this study, we investigated the strain-specific effect in variant calling of two major rice strains, Indica and Japonica. For this purpose, we performed variant calling with different types of samples and analyzed the predicted variants. We found that the number of variants called increased as the phylogenetically more different accessions from the reference were included in the samples. That is, the more the samples differ in their strains from the reference, the more variants are called. To understand the additionally called variants, we used the confusion matrix with associated metrics and showed that most of the additionally called variants were potential false positives. In addition, the predicted variants from samples of mixed strains produced potentially more false positives than from unmixed samples.

Based on these findings, we proposed a method to identify false positives and constructed the dbFP, which plays a role in the database of false positives. We applied the proposed method to dbSNP and verified its usefulness by showing the recovery of correct error rates. The characteristics of the analysis results were valid regardless of the coding region or the non-coding region. Although we discussed the results in detail for the case of the Japonica reference, the main results and their characteristics were also valid for the case of the Indica reference. These results support the robustness of the proposed method and its applicability as a post-process of variant calling. In this regard, this study provides recommendations for effective practices for strain-specific variant calling of species other than rice.

If the samples are not purebred, there is a certain probability that the variants called do not belong to the reference strain. To avoid this uncertainty, we used phylogenetically purebred samples to call actual (or true) variants. Although we used the set of actual variants as a benchmark for the correctness of the predicted variants, the actual variants identified in this study are by no means complete in the sense that the purebred samples used were not large enough. We expect the proposed method to become more effective and accurate as more purebred samples are used to call actual variants. Therefore, it would be desirable to have more purebred samples for actual variant calling so that a more reliable analysis can be performed. As a large database of actual variants becomes available, the false positives will be more accurately identified.

When large sample sets are available, we can independently repeat the same variant calling models with different sample sets. In such a case, we are able to report not only an estimate of performance metrics such as precision and recall, but also statistics related to these metrics. The statistics can provide additional information about the degree of accuracy of the proposed method in identifying false positives. In addition, an analysis similar to this study can be performed within the Indica strain. It is known through genomic studies that Indica can be further divided into two major strains of varietal groups, Indica I and Indica II [33]. An analysis similar to this study can provide new perspectives on the characteristics of these two Indica strains. These should be understood in future work.

## Acknowledgments

This work was supported by the National Research Foundation of Korea(NRF) grant funded by the Korean government(MSIT) No. 2022R1A4A1030348 (Y.-j. P. and C.-y. L.), No. 2021R1I1A3044289 (C.-y. L.), and by the research grant of the Kongju National University in 2022 (C.-y. L.).

## Code and script availability

We have released our analysis tool, a detailed script of the sampling procedure, input files, and output files so that anyone can reproduce the results. These are available at https://github.com/infoLab204/rice_strain.

## Supporting information

**S1 Fig.**
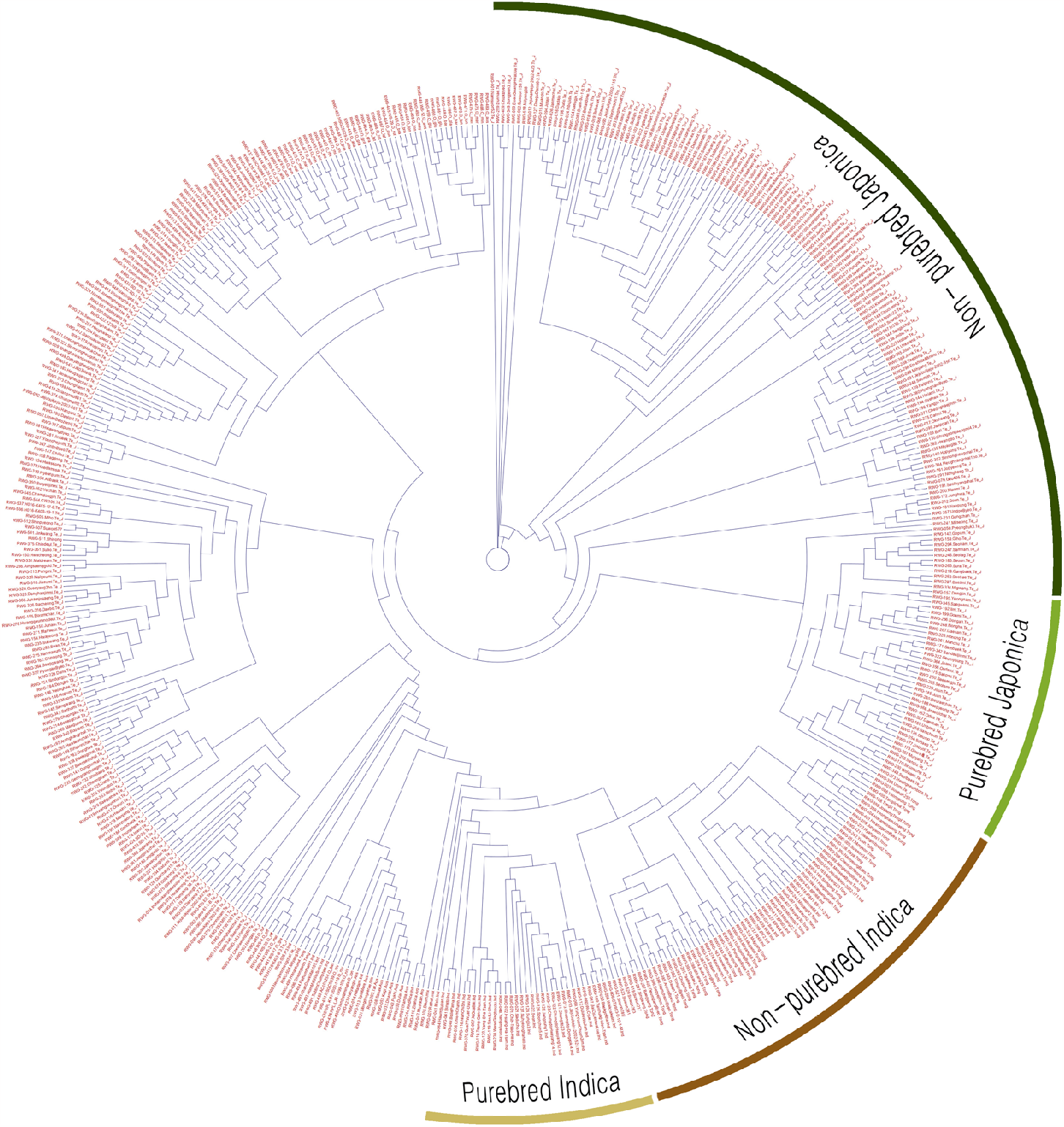
Phyloegentic tree. The phylogenetic tree of the core accessions in KRICE constructed using PHYLIP. The much higher resolution tree is available at https://github.com/infoLab204/rice_strain/blob/main/results/rice_circular.png.

**S2 Fig.**
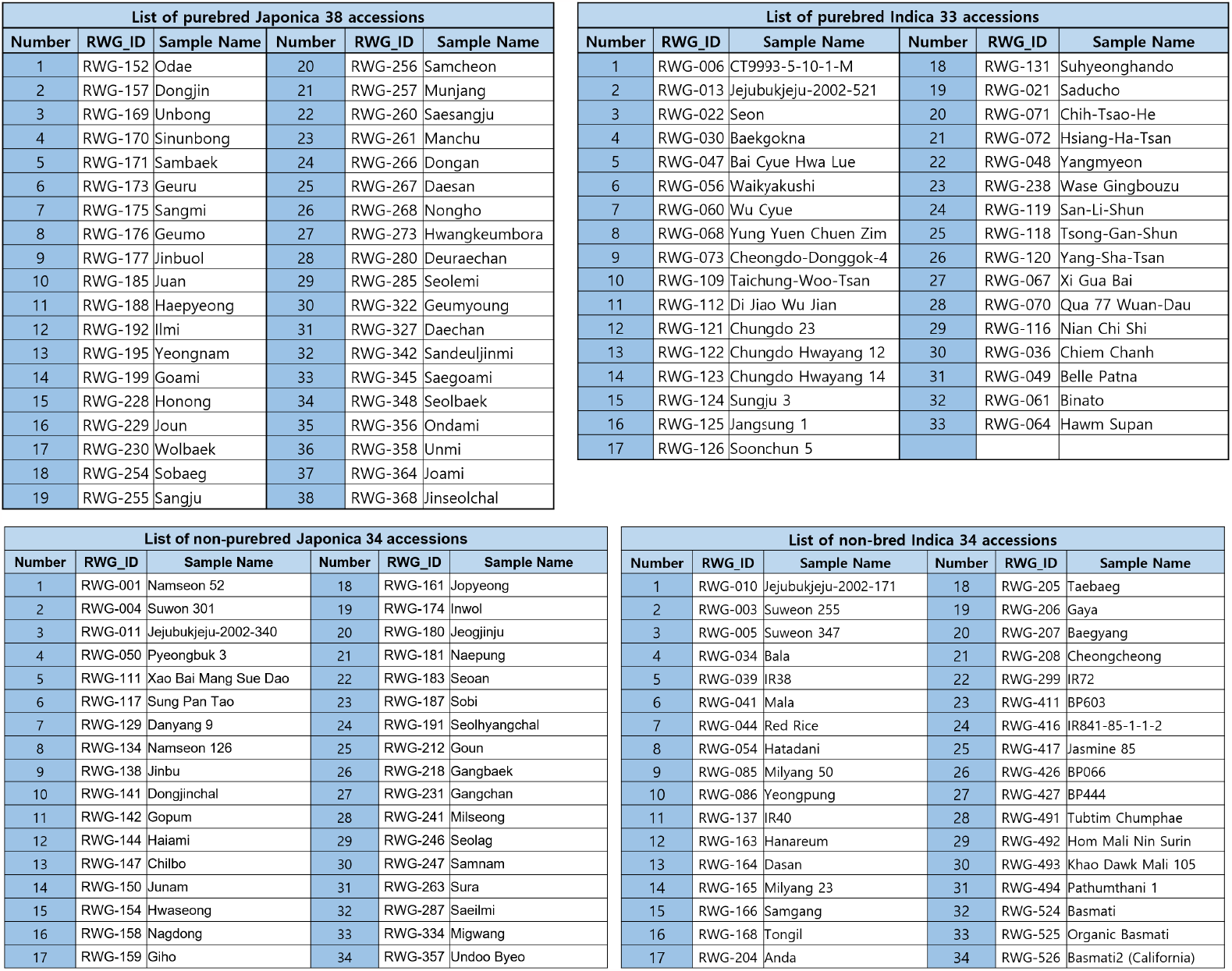
List of samples. (Top) The list of purebred Japonica and purebred Indica samples to call the actual variants. (Bottom) The list of non-purebred Japonica and non-purebred Indica samples to call the predicted variants.

**S3 Fig.**
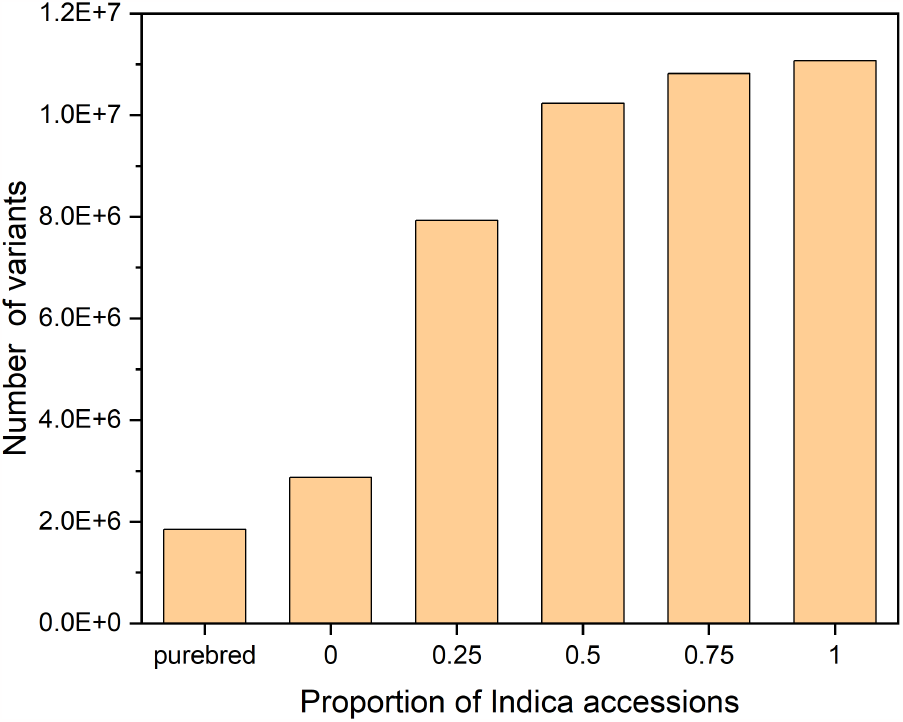
Variants in the non-coding region. The number of variants called from different models using the Japonica reference sequence in the non-coding region.

**S4 Fig.**
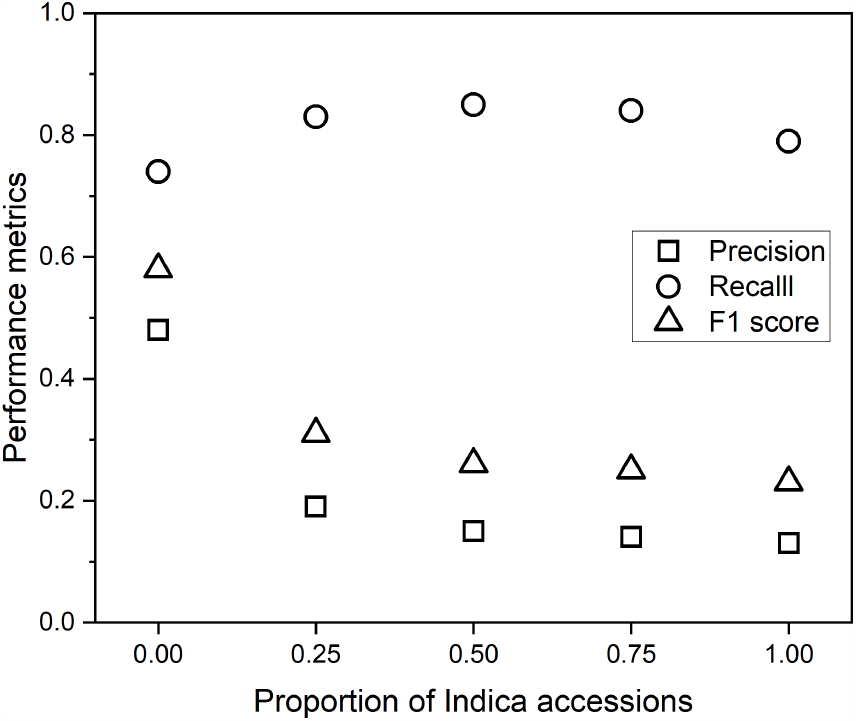
Metrics in the non-coding region. A plot of the performance metrics of precision, recall, and F1 score for different models in the non-coding region using the Japonica reference. Note that the horizontal axis represents the proportion of non-Japonica (i.e., Indica) accessions in the samples.

**S5 Fig.**
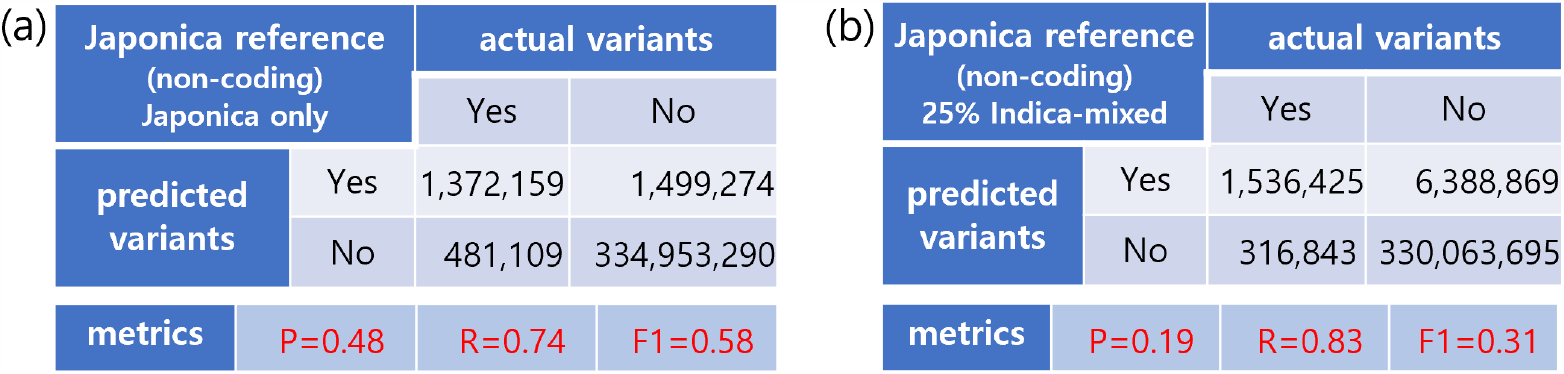
Confusion matrices in the non-coding region. Results of the confusion matrices and their metrics constructed from variant calling in the coding region using the Japonica reference from (a) the Japonica-only model and (b) the 25% Indica-mixed model.

**S6 Fig.**
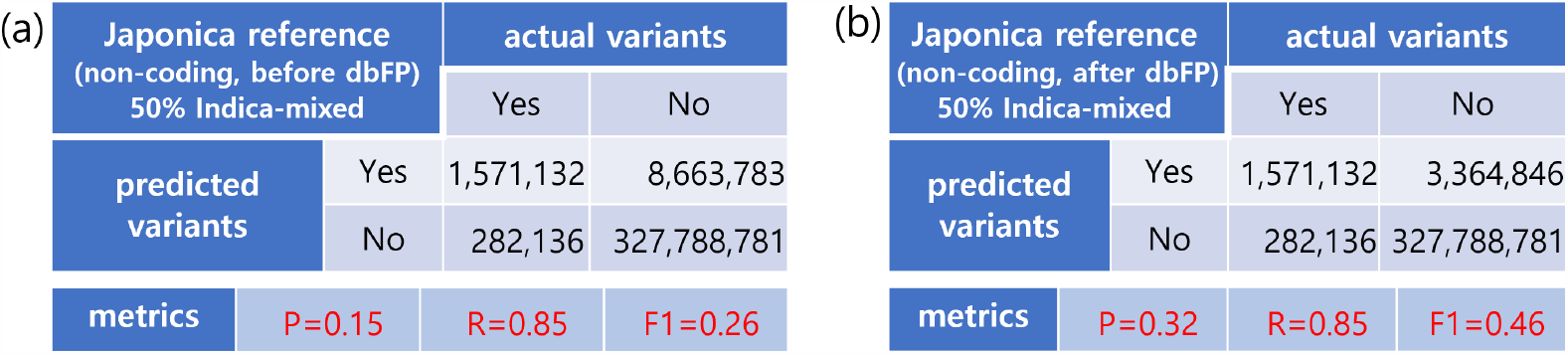
Before and after dbFP. Results of the confusion matrices and their metrics constructed from variant calling in the non-coding region with the Japonica reference using (a) the half-mixed samples from both strains and (b) the same as (a) except that the variants in the dbFP are excluded.

**S1 Table.** The number of false positives and precision. The number of false positives and precision from the models of different Indica-mixed proportions in the non-coding region.

| Model | 0% | 25% | 50% | 75% | 100% |
| --- | --- | --- | --- | --- | --- |
| Number of potential false positives | 1,499,274 | 6,388,869 | 8,663,783 | 9,257,316 | 9,604,602 |
| Precision | 0.48 | 0.19 | 0.15 | 0.14 | 0.13 |

**S1 File. Results of Indica reference**. The analysis results of the variant calling using the Indica reference sequence. The file for the results can also be found at https://github.com/infoLab204/rice_strain/blob/main/results/Indica_results.xlsx.

## Notes

### Competing Interest Statement

The authors have declared no competing interest.

